# A new humanized TCR transgenic mouse model to study citrullinated tenascin C reactive T cells relevant to rheumatoid arthritis

**DOI:** 10.64898/2026.02.12.705563

**Authors:** Marlene Schülein, Marcelo Afonso, Merel Sijbranda, Diego Velasquez Pulgarin, Anatoly Dubnovitsky, Monika Hansson, Bence Réthi, Fredrik Wermeling, Aaron Winkler, Lars Klareskog, Alexander Espinosa, Vivianne Malmström, Bruno Raposo

**Affiliations:** Department of Medicine, Division of Rheumatology, Karolinska Institutet, and Center for Molecular Medicine (CMM), Karolinska University Hospital, Stockholm, Sweden; Department of Hematology, St Jude Children’s Research Hospital, Memphis, TN 38105, USA; Inflammation and Immunology Research Unit, Pfizer Inc., Cambridge, MA, USA; Karolinska Center for Transgene Technologies (KCTT), Comparative Medicine, Karolinska Institutet, Stockholm, Sweden

## Abstract

**Objectives:** Autoreactive T cells recognizing citrullinated antigens, although rare and difficult to study, are implicated in rheumatoid arthritis (RA). To allow functional studies and manipulation of such T cells, we have generated and characterized transgenic mice expressing a TCR cloned from an RA patient and reactive with a prominent target, citrullinated tenascin C (citTNC).

**Methods:** A humanized TCR transgenic (hTCR-tg) mouse recognizing the citrullinated TNC22 (citTNC22) antigen in an HLA-DRB1*04:01 (HLA-DR4) restricted manner was developed by genetically engineering a chimeric TCR expressing murine constant domains and human V(D)J sequences. hTCR-tg mice were immune phenotyped in murine H-2^b^ and humanized HLA-DR4 backgrounds using full spectrum flow cytometry, cytokine ELISAs and fluorospot assays at steady-state and after antigen challenge. Additionally, we investigated the presence of antibodies to citTNC and arthritis following citTNC protein immunization.

**Results:** Thymic selection of hTCR T cells differed between the two MHC-II alleles, with a normal CD4+ T cell development observed solely under HLA-DR4 restriction. The chimeric hTCR maintained its citTNC22 specificity in vitro and ex vivo, without cross-reactivity to native TNC22 or other citrullinated autoantigens. hTCR-tg CD4+ T cells responded to antigen challenge in vivo and provided support to IgG class-switch and antigen-specific antibody production. Moreover, protein immunized hTCR-tg mice developed arthritis after periarticular challenge with citTNC.

**Conclusions:** hTCR-tg mice expressing an RA patient-derived autoreactive TCR develop functional antigen-specific and HLA-restricted CD4+ T cells. The combination of humanized HLA-DR4 and TNC22 mice will be a valuable tool in the development of antigen-specific therapies translatable to human disease.

## Introduction

MHC class II restricted T cell activation is central to the pathogenesis of the anti-citrullinated protein antibodies (ACPA)-positive subset of rheumatoid arthritis (RA)[1,2] and needed for the development of B cells/plasma cells producing the heavily somatically mutated antibodies reactive with citrullinated as well as other post-translationally modified targets[3]. Several known targets of ACPA have been shown to activate blood and synovial T cells from RA patients. These encompass both joint-specific (e.g. collagen type II or aggrecan[4–8]) and systemic antigens (e.g. fibrinogen, vimentin, alpha-enolase[8–12]). However, apart from candidate antigens based on these types of autoantibody reactivity, we do not have a full grasp of the breadth of T cell reactivities in RA, nor the contribution of each of these to the development of ACPA and RA itself. Additionally, we almost completely lack animal models where functions of such T cells can be investigated in detail, since current arthritis models that include antigen-specific immunity have so far mainly been based on immunities that are rarely seen in RA patients and are not specifically related to the disease-associated MHC class II gene variants. (e.g. CIA[13] and GPI-induced arthritis[14])

Post-translational modifications (PTM) of proteins have been shown to limit negative selection of developing thymocytes and consequently provide an explanation for the occurrence of autoreactive T cells in the naïve repertoire[15]. The identification of citrulline-reactive T cells in RA patients has therefore been a focus of research when trying to understand the aetiology of the disease and developing novel targets for personalized tolerance therapies. One of the T cell antigens receiving special attention in recent years is tenascin C (TNC)[16–20]. TNC is a large matrix protein poorly expressed at steady state but actively transcribed during inflammation (with pro-inflammatory properties itself) and tissue remodelling[21]. Stromal cells such as fibroblasts constitute the main producing cells of TNC [22]. Interestingly, CD4+ T cells recognizing distinct epitopes within citrullinated-TNC (citTNC) are found in peripheral blood and synovial fluid of RA patients[17,23], as well as in individuals at-risk of developing RA[19]. These observations indicate that autoreactive T cells targeting citrullinated antigens are present during disease and also develop prior to disease onset, implicating these cells in breach of tolerance mechanisms leading to RA. Importantly, citTNC-reactive T cells appear in higher frequency than all other autoreactive T cell specificities combined (e.g. citrullinated-fibrinogen, -vimentin or -alpha enolase)[17], suggesting a relevant role for this autoantigen in the development and/or progression of RA.

Recently, we have identified and cloned an expanded TCR expressed by an expanded clonal pool of T cells from the blood of a shared epitope positive (HLA-DRB1*04:01) RA patient recognizing the citrullinated TNC22 epitope (citTC22)[18]. A follow-up study assessing the interaction of this TCR with its cognate peptide-HLA (pHLA) complex demonstrates good TCR-pHLA affinity, with both citrulline motifs in the peptide being essential for a strong binding, whereas no recognition of the native peptide was observed in the context of HLA-DRB1*04:01(HLA-DR4)[20]. Making use of this well characterized autoreactive TCR, we have established a humanized T cell receptor transgenic (hTCR-tg) mouse strain whose CD4+ T cells recognize a citrullinated epitope within TNC (citTC22) in the context of HLA-DR4.

## Methods

### Generation of TNC22 hTCR-tg mice

The V(D)J sequences for a previously described RA patient-derived hTCR recognizing citrullinated peptide TNC22 (citTNC22)[18] were synthesized and inserted into TCR cassette vectors containing mouse constant TCRα or TCRβ sequences[24]. To ensure correct splicing and optimal expression, the V(D)J sequences were preceded by a human leader-intron sequence and followed by a splice donor and intronic sequence from mouse *Traj45* (for TCRα) or *Trbj1-2* (for TCRβ). The plasmids were linearized to remove the bacterial backbone and co-injected into pronuclei of C57BL/6J zygotes. Injected zygotes were implanted into pseudo-pregnant SWISS female mice. Pups were genotyped for chromosomal integration of the two plasmids, and positive founders were bred to wild-type C57BL/6J mice to generate chimeric hTCR-tg lines. Genotyping was done using the following primers: TCRα forward TGGCTATGGTACAAGCAGGA; TCRα reverse GTAGTCCCATCCCCAAAGGT; TCRβ forward AGAGACAGGAAGGCAGGTGA; TCRβ reverse CTGGCGCAGAAATACACAGA. Microinjections and embryo implantations were performed at Karolinska Center for Transgenic Technologies (KCTT). The resulting TNC22 hTCR-tg mouse strain was generated with approval by the Ethical Review Committee North, Stockholm County, and animals were handled in compliance with the guidelines at Karolinska Institutet.

### Mouse strains used

HLA-DR4 transgenic mice (B6.129S2-*H2-Ab1^tm1Gru^* Tg(HLA-DRA/H2-Ea,HLA-DRB1*0401/H2-Eb)1Kito) were purchased from Taconic Biosciences and bred with CD45.1 congenic C57BL/6J mice (JAX stock #002014; B6.SJL-*Ptprc^a^ Pepc^b^*/BoyJ) to establish wild type lines of recipient strains expressing either the murine H-2^b^ or the human HLA-DR4 allele (hereafter referred as WT.H-2^b^ or WT.DR4, respectively). hTCR-tg TNC22-reactive mice were backcrossed into a C57BL/6J background for ≥5 generations (TNC22.H-2^b^) and kept in a hemizygous fashion. TNC22.DR4 mice were generated by crossing TNC22.H-2^b^ mice with WT.DR4 mice, selecting for Ab1^-/-^ and CD45.2 expression. T cell deficient mice (JAX stock #002118; B6.129P2-*Tcrb^tm1Mom^*/J) were used to generate mice exclusively generating TNC22 hTCR-tg T cells (TNC22.TKO strain in a H-2^b^ background), as well as DR4 recipients deprived of polyclonal T cells (TKO.DR4 strain expressing the CD45.1 congenic allele). Littermates were used as control animals in every experiment. All animal experiments were performed with approval by the Ethical Review Committee North, Stockholm County, and animals were handled in compliance with the guidelines at Karolinska Institutet.

### Quantitative reverse-transcription PCR

RNA was extracted from sorted CD4+ and CD8+ T cells from TNC22.H-2^b^ and WT.H-2^b^ mice using the RNeasy Mini kit (Qiagen, cat# 74104), whereas cDNA was generated using the High-Capacity RNA-to-cDNA™ kit (Applied Biosystems, cat# 4387406) according to the manufacturer’s protocol. qPCR was performed for detection of the transgenic TCRβ (forward primer CCCTCCGGTGCAGTTCTTC; reverse primer GTGGAGTCACATTTCTCAGATCCT) and the housekeeping gene *Ppia* (forward primer CAGACGCCACTGTCGCTTT; reverse primer TGTCTTTGGAACTTTGTCTGCAA) and analysed using Bio-Rad CFX Maestro 2.3 (Version 5.3.022.1030).

### In vitro antigen stimulation

For determining antigen specificity and MHC-class II restriction of hTCR-tg CD4+ T cells, we established co-cultures of sorted CD4+ T cells and bone marrow-derived dendritic cells (BMDCs). Briefly, bone marrow from wild-type C57BL/6J mice (H-2^b^) or HLA-DR4 expressing mice (expressing solely HLA-DR4 or both MHC-II alleles – DR4.H-2^b^) was collected from their tibiae and femurs and meshed into a single cell suspension. After red blood cell lysis, cells were differentiated into BMDCs in DMEM culture medium containing 5% foetal calf serum, 100U penicillin plus 100ug/ml streptomycin and 1mM HEPES, supplemented with 100ng/ml murine Flt3l (Peprotech) and incubated for 8-10 days at 37°C, 5% CO_2_. After differentiation, BMDCs were evaluated for their expression of CD11c and high levels of MHC-II. 50-100 x 10^3^ CD11c^+^MHC-II^+/hi^ cells were used as antigen presenting cells per stimulatory condition, in a 96 well cell culture plate (Nunc). To collect T cells, secondary lymphoid organs [SLOs; spleen, inguinal lymph nodes (LN), axillary LN, brachial LNs, mesenteric LNs and cervical LNs] from naïve TNC22.H-2^b^ mice were harvested and meshed into a single cell suspension. T cells were negatively enriched using EasySep mouse T cell isolation kit (StemCell Technologies), followed by flow cytometry sorting of CD19-CD8-CD4+CD25- T cells (BD Influx or Sony SH800). 50-100 x10^3^ sorted CD4+CD25- T cells were cultured together with BMDCs and stimulated with native or citrullinated peptides (GenScript Biotech) at the indicated concentrations. The TNC22 peptide encompasses amino acids 1012-1026, with modifications at positions 1014 and 1016. Cell culture supernatant IL-2 and IFNγ, as well as IFNγ-producing cells were analysed by sandwich ELISA and fluorospot (MabTech), respectively (see below for details).

For in vitro recall responses, single cell suspensions of spleens and/or draining lymph nodes were prepared in RPMI medium supplemented with 5% foetal calf serum, 100U penicillin plus 100ug/ml streptomycin, 1mM HEPES and 50uM β-mercaptoethanol. 0.5-1 x 10^6^ splenocytes or draining lymph node cells were stimulated with 5-10ug/ml of desired peptides. Unstimulated conditions as well as plate-bound anti-CD3 (2ug/ml, clone 145-2C11, BD Pharmingen) and soluble anti-CD28 (1ug/ml, clone 37.51, Biolegend) conditions were always included to assure a negative (background) and positive control culture condition.

### Cell transfer experiments, immunizations

For cell transfer experiments, SLOs from naïve TNC22.TKO mice (containing solely hTCR-tg T cells) were harvested and prepared for flow cytometry cell sorting as described above. 1-5 x 10^5^ sorted CD4+CD25- T cells were transferred to TKO.DR4 recipient mice depending on the experimental setting. Transferred T cells were left to expand in TKO.DR4 recipients for a week, after which recipient mice were immunized with 50ug citTNC22 peptide in 100ul of complete Freund’s adjuvant (CFA), in a 1:1 emulsion. Spleens and inguinal lymph nodes were collected for analysis 13 days post immunization.

To investigate the general polyreactivity to citTNC22 by HLA-DR4 transgenic mice, WT.DR4 animals were immunized at the base of the tail with 50ug citTNC22 peptide in 100ul of CFA, in a 1:1 emulsion and boosted 21 days later with 25ug peptide in 50ul incomplete Freund’s adjuvant (IFA), in a 1:1 emulsion. Blood samples were collected at days 21 and 52 post initial immunization.

To assess antibody responses to TNC mediated by hTCR-tg T cells, TKO.DR4 mice receiving CD4+CD25- T cells from either TNC22.TKO or WT.H-2^b^ mice were primed (day 0) and boosted (day 14) with 50ug of a citrullinated fragment of tenascin C (citTNC-T4), together with 5uM CpG-B (ODN-1668, InVivoGen) in 100ul of a saline solution at the base of the tail. Plasma was collected at days 14 and 35 and assessed for IgM and IgG titres against native- and citrullinated-TNC-T4 protein (natTNC-T4 and citTNC-T4, respectively).

### Cytokine ELISA and Fluorospot

A sandwich ELISA was used to detect IL-2 and IFNγ production in response to native and citrullinated peptides after in vitro co-cultures of T cells-BMDCs, as well as ex vivo restimulation of T cells. 96 well MAXISORP plates (Nunc) were coated with 2ug/ml anti-IL2 (clone JES6-1A12) or anti-IFNγ (clone R4-6A2) overnight at 4°C, followed by blocking with 2% BSA in PBS. Plates were washed with PBS-Tween (0.05%) between every step. After blocking, cell culture supernatants were added to the plate and incubated for a minimum of 2h at room temperature or overnight at 4°C. Detection was done with 0.5ug/ml biotinylated anti-IL-2 (clone JES6-5H4) or anti-IFNγ (clone XMG1.2) followed by streptavidin-HRP. ELISAs were developed using TMB substrate (Biolegend) and the reaction stopped with 0.5M H_2_SO_4_ after 10min. ELISA data is represented in pg/ml or absorbance values, the later used when all samples and respective controls could be analysed in one single 96 well ELISA plate.

IFN-γ producing cells were detected using mouse IFNγ Fluorospot assays (MabTech), according to the manufacturer’s protocol and analysed using an IRIS 2 Fluorospot reader (MabTech). Data presented shows the number of IFN-γ producing cells per million cells plated, or the average detected intensity of the signal per cell (RSV550).

### ELISAs for antibody titres

For antigen-specific antibody titre detection, ELISA plates were coated with 5ug/ml natTNC-T4 or citTNC-T4 overnight at 4°C, followed by blocking with 2% BSA in PBS. Plates were washed with PBS-Tween (0.05%) between every step. After blocking, plasma samples were added to the plate at a predetermined dilution in RIA buffer (10mM Tris-HCl, 350mM NaCl, 1% tween-20, 10% SDS, 1% BSA, pH=7.6), or a series of dilutions, and incubated for a minimum of 2h at room temperature or overnight at 4°C. Goat F(ab’)2 anti-mouse IgG HRP-conjugated (Jackson Immuno Research, cat# 115-035-062) were used as secondary antibodies for the detection of plasma IgG. ELISAs were developed using TMB substrate (Biolegend) and the reaction stopped with 0.5M H_2_SO_4_ after 10min. Data is represented as absorbance values (used when all samples and respective controls could be analysed in one single 96 well ELISA plate) or arbitrary units (where a positive control pooled sera sample was added to each ELISA plate to generate a standard curve and extrapolate the unknown plasma reactivities).

The commercial anti-CCP2 ELISA kit (Immunoscan CCPlus; SVAR Life Science AB) was used to quantify the anti-citrulline specific antibody reactivity. A goat F(ab’)2 fragment anti-mouse IgG + IgM HRP conjugated (Jackson Immuno Research, cat# 115-036-068) was used as secondary antibody.

### Flow cytometry analysis

For flow cytometry analysis, fresh blood, thymi or SLOs were collected from mice and organs meshed through a 40um nylon mesh into a single cell suspension. After red blood cell lysis, cells were treated with Fc block (anti-CD16/CD32) and viability dye (Zombie NIR, Biolegend) in PBS for 20 minutes at 4°C. To minimize antibody aggregation and interference, surface staining was split into two sequential staining steps. Cells were first stained with a multicolour antibody cocktail (cocktail #1) for surface staining (suppl. table 1) in PBS containing 0.5% FBS and 2mM EDTA for 1 hour at 4°C, followed by a 16-hour overnight staining with multicolour antibody cocktail (cocktail #2; suppl. table 2). For intracellular staining, cells were fixed and permeabilized with Foxp3/transcription factor fixation buffer (eBioscience) for 30min at 4°C, followed by staining with AlexaFluor 647-Foxp3 antibody for 30 minutes at 4°C. Stained cells were resuspended in PBS containing 0.5% FBS, 2mM EDTA and acquired in a 5-Laser Cytek Aurora flow cytometer (Cytek BioSciences). Data was analysed using FlowJo v10 (BD Biosciences). A cocktail of anti-TRBV antibodies was used to detect the highest possible frequency of polyclonal T cells in both wild-type and hTCR-tg littermates (WT.H-2^b^ and TNC22.H-2^b^, as well as WT.DR4 and TNC22.DR4; suppl. table 1). This mixture of anti-TRBV antibodies is limited to what is commercially available and can detect ≈50% of all polyclonal T cells, allowing us to extrapolate the frequency of hTCR-tg T cells in a mixed T cell population.

#### In vitro citrullination of proteins

Due to its large molecular weight (≈241kDa; Uniprot entry P24821), human TNC was re-expressed in E. coli in different fragments of 30-45kDa. Fragment T4 (TNC-T4) encompasses the amino acid sequence 981-1270 and contains the TNC22 peptide antigen. Polypeptide including amino acids 981-1270 of unprocessed tenascin C (numbering includes signal peptide) followed by a short linker and C-terminal His6-tag (LEHHHHHH) was overexpressed from pET24 vector in *E. coli* BL21(DE3) STAR cells. Protein was expressed insoluble in inclusion bodies. Expression was induced by addition of 1 mM IPTG for 4 h at 37°C. Cells were harvested by 10min centrifugation at 5,000g at 4°C, resuspended in 20 mM Tris-HCl pH 8 and frozen at - 20°C. After thawing, DNAse, 0.5mM MgCl, 1mM MnCl, pefablock and complete protease inhibitors cocktail (Roche) were added to the cell suspension. Cells were disrupted by sonication and inclusion bodies were harvested by 20min centrifugation at 10,000g. Inclusion bodies were washed 5x with 20mM Tris-HCl pH 8, 0.5mM MgCl, 1mM MnCl, DNAse, once with 0.05% Triton X100 in 20mM TRIS-HCl pH 8 and finally, 2x with distilled water. Purified inclusion bodies were solubilized in 8M urea, 20mM Tris-HCl pH 8.0 and purified using IMAC chromatography under denaturing conditions in 6M Urea. To remove LPS, the pH of the solution was titrated down to 3.7 and protein was loaded onto 5mL HiTrap™ Q HP column (Cytiva). The flow-through was collected and directly loaded onto 5mL HiTrap™ SP HP column (Cytiva). Protein was eluted in 6M Urea, 20mM Tris-HCl, 0.5M NaCl, concentrated on 5 kDa cut-off Vivaspin and purified by size-exclusion chromatography using HiLoad™ 16/600 Superdex™ 200 PG column (Cytiva) in 6M Urea. Before in vitro citrullination, urea was removed by dialysis against 100mM Tris-HCl pH 7.5.

Citrullination of the TNC-T4 protein fragment was performed in 100mM Tris-HCl containing 50mM CaCl2, 5mM DTT and incubated overnight at 37°C with 1U/mg PAD2 (rabbit muscle; Merck-Sigma cat# P1584). Human fibrinogen (Merck-Sigma) was PAD2 treated the same way at the same time as a positive control protein. After incubation, 20mM EDTA was added to the protein solution, followed by dialysis to PBS. Citrullination of the arginine motifs within the TNC22 peptide was confirmed by cellular assay using TNC22 hTCR-tg CD4+ T cells (fig. 5A), whereas the general citrullination efficiency was confirmed by western blot using the co-citrullinated fibrinogen sample. Briefly, protein samples were denatured in 4X NuPAGE LDS Sample Buffer (Thermofisher cat. NP0007) at 70°C for 15 minutes and separated with SDS-PAGE using Bolt™ Bis-Tris 4-12% gel (Thermofisher). Proteins were transferred onto a PVDF membrane (Invitrogen, cat# LC2002) at 25V for 2 hours in transfer buffer containing 10% MeOH. The membrane was blocked with 3% BSA in PBS and incubated overnight with 1ug/ml of primary antibodies (ACPA clone 1325:01B09[25]; suppl. fig. 4B). Detection was done with a HRP conjugated secondary anti-mouse IgG (1:5000; Jackson Immuno Research, cat# 115-036-071) for 1h in 1% BSA in PBS, followed by Clarity Western ECL Substrate (BioRad cat# 170-5060).

### Immunofluorescence staining of TNC

Arthritic hind-paws from animals recently undergoing collagen antibody-induced arthritis (CAIA) were collected at day 14 post disease induction (for detailed protocol of disease induction see ref. [26]). Freshly collected paws were fixed in 4% paraformaldehyde overnight and decalcified in 0.5M EDTA for three weeks. Tissue was then embedded in OCT Cryomount (Histolab Products AB, Sweden) and sectioned in 10um slides. Sections were incubated with glycine and subsequently with PBS 0.1% saponin. Afterwards, slides were stained using biotinylated monoclonal mouse anti-collagen type II (clone M21139, Absolute Antibody, Catalog # Ab02802-3.0, 10 mg/ml) and monoclonal rabbit anti-tenascin C (clone EPR4219, Abcam, Catalog # ab108930, 4.85 mg/ml) in PBS 0.1% saponin supplemented with 3% mouse serum. After overnight incubation at room temperature, slides were washed and blocked with PBS 0.1% saponin supplemented with 3% mouse serum and 1% goat serum. Subsequently, slides were stained with Streptavidin-Alexa Fluor 647 (ThermoFisher, Catalog# S21374, 1:1,000) and goat-anti-rabbit Cy3 (Jackson ImmunoResearch, Catalog# 111-166-046, 1:400) secondary antibodies and Hoechst 33342 DNA dye (ThermoFischer, Catalog# 3570, 1:50,000) in PBS 0.1% saponin supplemented with 3% mouse serum. Images were acquired on a LSM 880 confocal microscope without Airyscan (Zeiss).

### Arthritis induction

As described in the cell transfer paragraph above, CD4+CD25- T cells were sorted from SLOs of TNC22.TKO or WT.H-2^b^ mice. 2×10^6^ sorted cells were stimulated in vitro for 3 days with 5ug/ml of citTNC22 peptide, in the presence of TKO.DR4 splenocytes. After stimulation, cells were washed in cold PBS and transferred i.p. to recipient TKO.DR4 mice (n=4). Recipient mice were immunized the day after with a mix of 25ug citTNC22 peptide and 50ug citTNC-T4 protein in 100ul CFA emulsion (1:1). Ten days later, mice were periarticularly s.c. challenged with 50ul citTNC22 peptide and citTNC-T4 protein in the right hind ankle, while the left hind ankle served as a control. Inflammatory arthritis was assessed both qualitatively (degree of swealing: 0 - no arthritis, 1 – mild arthritis in ankle or foot, 2 – mild arthritis in both ankle and foot, 3 – severe arthritis in ankle or foot, 4 – severe arthritis in ankle/foot and mild arthritis in foot/ankle, 5 – severe arthritis in both ankle and foot, 6 – severe arthritis in both ankle, foot and digits) and quantitatively using a calliper to measure the thickness of the ankle joint and swelling of the foot above the knuckles.

### Statistical analysis

When comparing two groups the Mann-Whitney test was used for samples with n<10, otherwise an unpaired t-test was used for samples with n≥10. One-way ANOVA with Holm-Šídák’s multiple comparisons test or Friedman test with Dunn’s multiple comparison’s test were used where appropriate. For analysis of paired data, paired t-test or Wilcoxon matched-pairs signed rank test was used. Statistical significance of arthritis severity was assessed with a 2way ANOVA with Tukey’s multiple comparison test. P<0.05 was considered significant. Statistical analysis was performed using GraphPad Prism Version 10.3.0.

## Results

### DR4 transgenic mice respond to citTNC22 peptide

Recent work by McElwee and colleagues showed that the frequency of T cells recognizing citrullinated (auto)antigens in DR4 transgenic mice is dependent on the antigen, indicating different levels of immune tolerance to the various PTM antigens[27]. To extend on this, we evaluated the T cell response to one of the TNC peptides previously identified as a T cell epitope in RA patients[17–19] (fig. 1A). We observed that several DR4 mice challenged with the antigen mounted an immune response targeting the citrullinated peptide (citTNC22), with some reactivity also to its native version (natTNC22; fig. 1B-C). Additionally, a B cell response towards citrullinated antigens was also observed in these animals, measured by the clinically used CCP2 test, initially modest and with increasing titres after antigen boosting (fig. 1D). Overall, the polyclonal T cell response observed in DR4 mice indicates that this HLA-restricted PTM autoantigen can be studied in an experimental murine setting. At the same time, the variability in immune responses demonstrate that studies looking at citrulline-specific T cell reactivities would benefit from a more robust and reproducible model, where antigen-specific T cells can be easily monitored.

**Figure 1.**
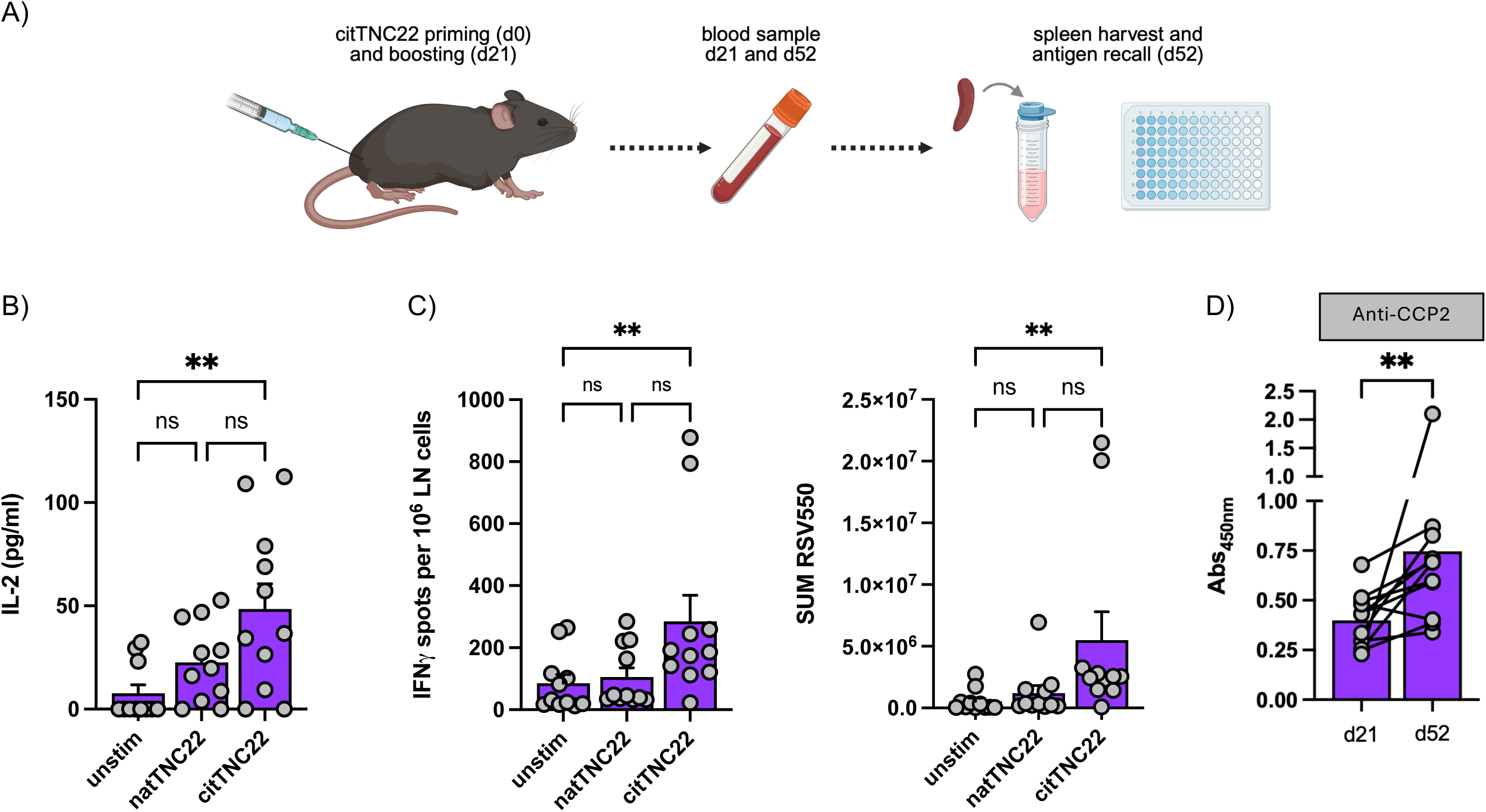
Polyclonal T cell reactivity to citTNC22 in DR4 mice. A) Schematic representation of the experimental setup. B-C) Supernatant IL-2 levels (B) and IFNγ producing cells (C) from in vitro stimulated splenocytes collected 52 days post initial immunization. Friedman test with Dunn’s multiple comparison’s test. ** - p<0.01; ns – not significant D) Anti-CCP antibody response in the serum of citTNC22-immunized mice at days 21 and 52 post immunization (1:25 dilution). Wilcoxon matched-pairs signed rank test. ** - p<0.01

### Flow cytometry characterization of hTCR-tg T cells

To increase the translational aspect of RA-related T cell tolerance and autoreactivity efforts, we made use of our previous TCR repertoire data set from citTNC22 reactive T cells from DR4+ RA patients[18] and generated a TCR transgenic mouse expressing a RA-patient derived paired alpha beta TCR sequence with high affinity for TNC22 presented by DR4[20]. This novel mouse strain (from here on referred to as TNC22 mice) was developed in a C57BL/6J background (TNC22.H-2^b^) using a well-established construct[24] to express a murine/human hybrid TCR (suppl. fig. 1A-B).

Phenotypic analysis of TNC22.H-2^b^ mice using flow cytometry revealed differences in thymic T cell development compared to WT.H-2^b^ littermates in line with observations from other TCR transgenic settings[24,28,29]. Here, TNC22.H-2^b^ mice displayed a decreased double positive CD4+CD8+ T cell population as well as decreased single positive CD4+ T cell compartment (fig. 2A), indicating a possibly defective positive selection of the hTCR. Double negative thymocytes were primarily observed within the final stage of TCR rearrangement (DN4), suggesting an advantageous T cell development and existence of a pre-TCRalpha, as previously described[29,30] (fig. 2B). Development of hTCR-tg T cells was further confirmed by the expression of transgenic mRNA transcripts both in the thymus and peripheral T cells in these mice (suppl. fig. 2A-B).

**Figure 2.**
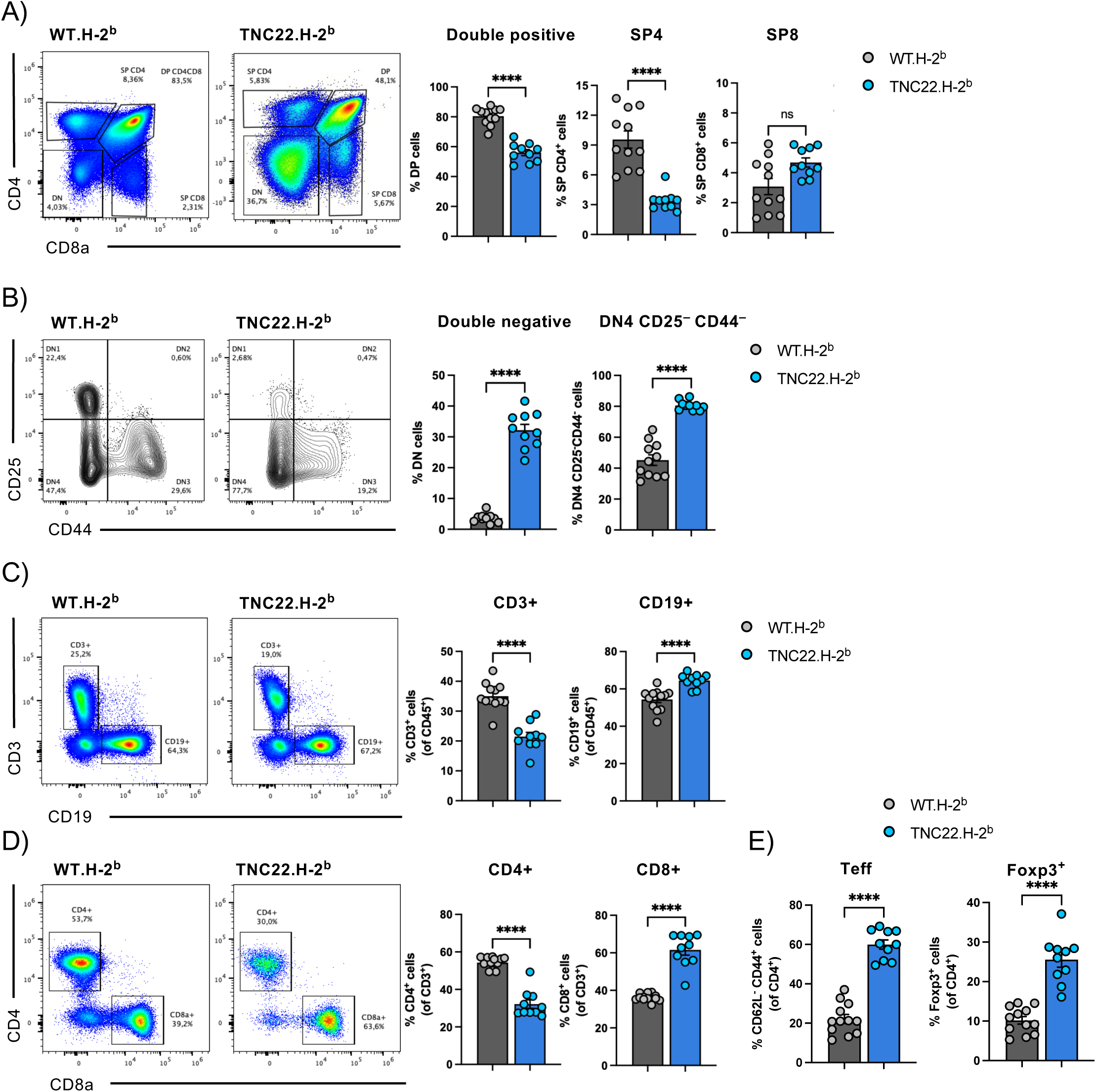
Thymic development and peripheral characteristics of TNC22 hTCR-tg T cells in H-2^b^ background. A) Flow cytometry plots of CD4 vs. CD8a and frequencies of thymic double positive (DP), single positive CD4^+^ (SP4) and CD8^+^ (SP8) cells. B) Flow cytometry plots of double negative (DN) stages and frequencies of DN and DN4 T cell populations in the thymus. C) Flow cytometry plots of CD3^+^ vs. CD19^+^ splenic cells and frequencies of these populations. D-E) Flow cytometry plots of splenic CD4^+^ vs. CD8^+^ and frequencies of these populations (D), as well as frequencies of effector CD4^+^ T cells (CD62L^-^CD44^+^) and Foxp3^+^ cells (E). Mann-Whitney test. **** - p<0.0001; ns – not significant.

The differences in thymic T cell development were also reflected in the periphery of TNC22.H-2^b^ mice, with the reduction of thymocyte count affecting the peripheral immune cell count (suppl. fig. 2D-E), particularly in terms of lower splenic T-cell frequencies (fig. 2C), as well as altered CD4:CD8 ratios (≈1.5 fold in WT.H-2^b^ and ≈0.5 fold in TNC22.H-2^b^; fig. 2D). Using a cocktail of anti-TRBV antibodies to identify murine polyclonal T cells (suppl. table 1), we could estimate a ≈50% frequency of TNC22-specific T cells in TNC22.H-2^b^ mice (suppl. fig. 2F-G), supporting that central T-cell selection is heavily biased to the transgenic hTCR. For comparison, the widely used OT-II TCR transgenic mice express the transgenic TRAV2 and TRBV5 alleles in >90% of circulating T cells[28]. Interestingly, we observed an increased frequency of activated T cells (CD62L-CD44+) as well as Foxp3+ regulatory T cells in spleens of TNC22.H-2^b^ mice compared to their WT.H-2^b^ littermates (≈22% to ≈60% CD62L-CD44+ cells and ≈10% to 25.5% Foxp3+ cells, respectively; fig. 2E). This observation strongly suggests that the cloned hTCR recognizing the citTNC22 peptide in the context of HLA-DR4 may also recognize another (yet unknown) antigen at steady state in the context of the murine H-2^b^ allele. This unknown antigen is likely to be present at high frequency, given the high levels of peripheral CD4+Foxp3+ T cells, as well as thymus-derived regulatory T cells (suppl. fig. 3A). Although we cannot completely differentiate polyclonal from transgenic T cells due to lack of specific antibodies for the human Vβ allele (transgenic T cells were estimated to be half of the CD4+ T cell population in TNC22.H-2^b^ mice), these phenotypes are likely associated with the expression of the transgenic hTCR.

### hTCR-tg CD4+ T cells are antigen-specific and HLA-restricted

The human HLA-DRB1*04:01 and murine H-2^b^ molecules have considerable differences in the amino acid composition of their peptide binding pockets[31,32]. Hence, it is very unlikely that the T cell phenotypes observed at steady state in TNC22.H-2^b^ mice, in particular the elevated frequencies of effector (Teff; CD62L-CD44+) and regulatory (Foxp3+) CD4+ T cells, could be due to recognition of the same antigen (i.e. citTNC22) in the context of both MHC-II alleles. In fact, TNC is poorly expressed under steady state conditions[21], and therefore unlikely to cause such a strong effector and regulatory T cell imprinting in these animals.

To confirm the antigen specificity and functionality of the hTCR-tg T cells without an eventual influence of the occurring Treg cells, we used the expression of CD25 as a proxy for Foxp3+ T cells and used it to sort non-Treg cells in our continued studies. Additionally, we crossed the hTCR-tg mice to a TCRβ knockout background (TNC22.TKO) to eliminate the influence of polyclonal T cells in our readouts. Flow cytometry sorted peripheral CD4+CD25- T cells from TNC22.TKO mice were co-cultured with bone marrow derived dendritic cells (BMDCs) from mice expressing either the murine H-2^b^ allele alone or a combination of both human HLA-DR4 and murine H-2 ^b^ (DR4.H-2^b^; fig. 3A). Peptide stimulation with either the native (natTNC22) or the citrullinated TNC22 (citTNC22) peptide confirmed that hTCR-tg CD4+ T cells recognize their cognate citTNC22 antigen solely in the context of human HLA-DR4, while showing no reactivity to this antigen when presented by murine MHC-II H-2^b^-expressing antigen presenting cells (fig. 3B). Additionally, hTCR-tg T cells showed no response to the natTNC22 peptide (fig. 3B) or other citrullinated antigens whose T cell reactivities are often found in RA patients (suppl. fig. 4A). Together, this data clearly indicates that hTCR-tg T cells respond to their cognate antigen in an HLA-DR4 restricted manner, and that the antigen leading to the T cell selection phenotypes showed earlier (fig. 2E and suppl. fig. 3A) in the context of H-2^b^ are unrelated to TNC22.

**Figure 3.**
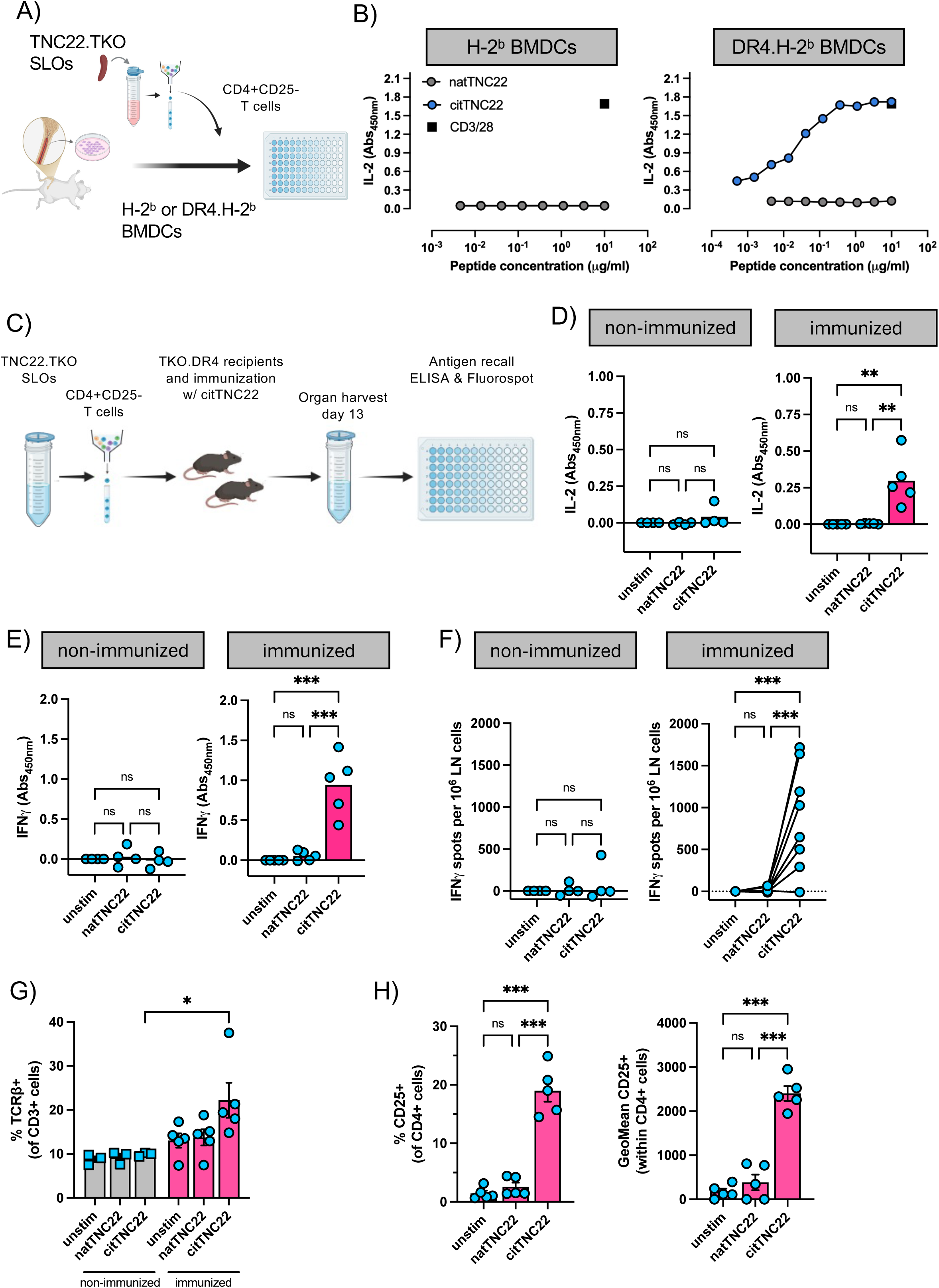
Reactivity of TNC22 transgenic T cells in vitro and ex vivo. A) Schematic representation of TNC22 transgenic CD4^+^CD25^-^ T cell sorting and co-culture with H-2^b^ or DR4.H-2^b^ BMDCs. B) Supernatant IL-2 levels secreted by TNC22 transgenic CD4^+^CD25^-^ T cells when stimulated by H-2^b^ BMDC or DR4.H-2^b^ BMDCs. C) Schematic representation of TNC22 CD4^+^CD25^-^ T cell sorting and transfer into TKO.DR4 recipient mice, followed by immunization and antigen recall. D-F) Supernatant IL2- (D) and IFNγ (E) by splenocytes after antigen recall, or IFNγ producing lymph node cells after antigen stimulation in vitro. G-H) Frequencies of TCRβ+ cells (G), CD4^+^CD25+ T cells and geometric mean (GeoMean) of CD25 (H) in CD4^+^ T cells from TKO.DR4 recipient mice after antigen recall. One-way ANOVA with Holm-Šídák’s multiple comparison’s test. * - p<0.05, ** - p<0.01 and *** - p<0.001; ns – not significant.

To further assess the functionality of the hTCR-tg T cells in vivo, flow cytometry sorted peripheral CD4+CD25- hTCR-tg T cells from TNC22.TKO mice were transferred into TKO.DR4 recipient mice. These mice express the human HLA-DR4 instead of the murine H-2^b^ molecule and do not have endogenous αβT cells, being the CD3+ T cell compartment exclusively composed of γδT cells. Recipient mice were either left untouched (non-immunized) or were immunized with citTNC22 peptide in CFA one week after cell transfer and spleens and draining lymph nodes collected from all recipient mice 13 days post immunization to assess antigen recall responses (fig. 3C). Antigen recall showed T-cell reactivity to the citrullinated but not the native peptide, confirming that hTCR-tg T cells were activated in vivo by their cognate antigen when presented on HLA-DR4 (fig. 3D-I). T cell recipient mice that did not get immunized showed no antigen recall response and no expansion of CD3+TCRβ+ T cells as compared to immunized mice, further confirming the antigen-specific activation of the hTCR-tg T cells in the context of HLA-DR4 (fig. 3D-G and suppl. fig. 5A).

### Flow cytometry characterization of hTCR-tg T cells in the context of HLA-DR4

To start understanding the developmental cues and characteristics of citrulline autoreactive T cells in a relevant MHC-II, TNC22.H-2^b^ mice were crossbred with DR4 mice to create double transgenic mice carrying both the hTCR and the human HLA-DR4 (TNC22.DR4), but without the murine H-2^b^ allele. Alike in the murine H-2^b^ background, flow cytometry analysis of thymocytes evidenced a decrease in frequency of double positive T cells and increase of double negative T cells in TNC22.DR4 mice compared to WT.DR4 littermates (fig. 4A-B). Likewise, most double negative T cells were in the final stages of TCR rearrangement, indicating expression of the transgenic hTCR (fig. 4B). Similarly to what has been described in other TCR transgenic systems[14], the co-expression of the transgenic TCR and its selective HLA allele did not promote skewing of mature CD4+ T cells, either in the thymus or periphery. The similar frequencies of double positive thymocytes in the “selecting” HLA-DR4 and “non-selecting” H-2^b^ backgrounds (fig. 4A and 2A, respectively) indicate that the transgenic TCR might be poorly selected in both cases. Nevertheless, phenotypic analysis of peripheral lymphocytes showed significant differences between the two MHC-II backgrounds. Unlike TNC22.H-2^b^ mice, TNC22.DR4 mice display a higher frequency of total T cells in the periphery, while the B cell levels were not significantly affected (fig. 4C). Interestingly, whereas the frequency of peripheral CD4+ T cells was strongly reduced in TNC22.DR4 mice, this reduction was not accompanied by an increase in the frequency of CD8+ T cells. Instead, TNC22.DR4 mice display a significantly increased CD4-CD8- double negative T cell population (fig. 4D), a phenotype previously described as resulting from peripheral antigen encounter[33]. Whether the antigen being recognized and leading to the eventual downregulation of CD4 and/or CD8 after antigen encounter is citTNC22 or another unrelated antigen (e.g. microbiota-derived or murine-restricted) presented by HLA-DR4 or the murine MHC-I (H-2K^b^/-2D^b^), it remains to be elucidated.

**Figure 4.**
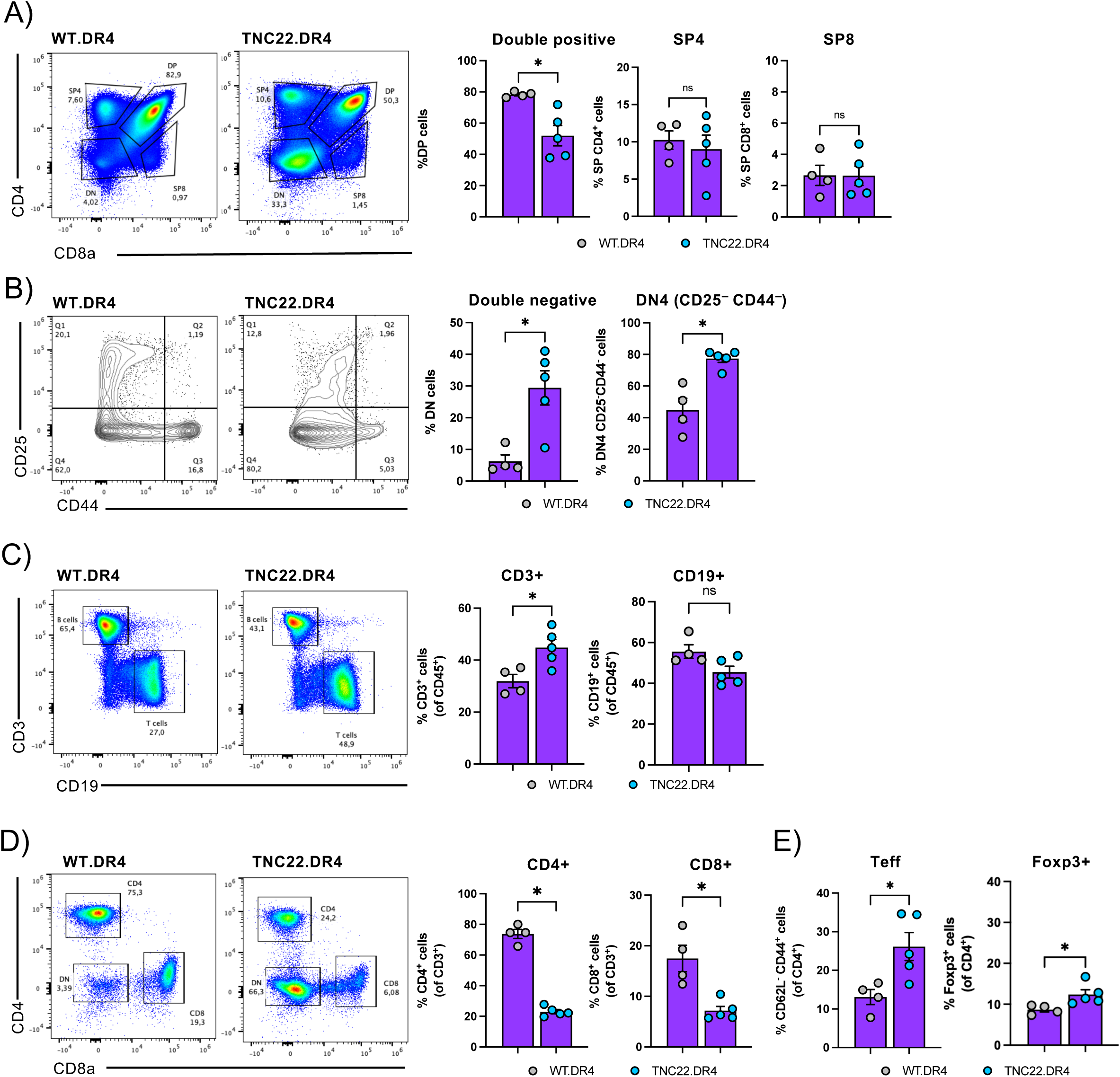
Thymic development and peripheral characteristics of citTNC22-reactive T cells in DR4 mice. A) FACS plots of CD4^+^ vs CD8^+^ and frequencies of double positive (DP), single positive CD4^+^ (SP4) and CD8^+^ (SP8) cells. B) FACS plots of double negative (DN) stages and frequencies of DN and DN4 populations in the thymus. C) FACS plots of CD3^+^ vs CD19^+^ splenic cells and frequencies of these populations. D-E) FACS plots of splenic CD4^+^ vs. CD8^+^ and frequencies of these populations (D), as well as frequencies of effector CD4^+^ T cells (CD62L^-^CD44^+^) and Foxp3^+^ cells (E). Mann-Whitney test. * - p<0.05; ns – not significant.

Importantly, the development of hTCR-tg T cells in the DR4 background resulted in an increased frequency of αβT cells as well as increased expression of the transgenic hTCR to ≈80% (suppl. fig. 6D-E; anti-TRBV antibody cocktail detects half of all polyclonal T cells, which in TNC22.DR4 mice it regards to ≈10% of all αβT cells), compared to ≈50% in the murine H-2^b^ background (suppl. fig. 2F-G). Peripheral hTCR-tg T cells from TNC22.DR4 mice display an activated effector phenotype (CD62L-CD44+) and increased Foxp3+ Treg population (fig. 4E), as observed in TNC22.H-2^b^ mice (fig. 2E). However, both T cell compartments were significantly reduced in TNC22.DR4 mice in comparison to the latter, demonstrating that the HLA-DR4 expression partially rescues the differentiation and activation status of these cells at steady state. This observation was consistent between secondary lymphoid organs draining peripheral or mucosal tissues (spleen, fig. 2C-E and 4C-E; inguinal and mesenteric lymph nodes [iLN and mLN], suppl. fig. 3B-C and 6B-C).

### hTCR-tg T cells support B cell class-switch and anti-citrulline antibody responses

One of the standing questions in the field of RA is the origin of the immune response leading to ACPAs and their contribution to arthritis development. Hence, we next sought to explore whether the hTCR-tg T cells are able of supporting B cell activation and antigen-specific antibody production.

The initial identification of citTNC22-reactive T cells in RA patients was based on in silico prediction of peptides able to dock the human HLA-DR4 molecule[17]. To confirm the biological processing and presentation of this peptide by antigen-presenting cells, we in vitro citrullinated a protein fragment of TNC that contains the TNC22 epitope (citTNC-T4). When co-culturing hTCR-tg CD4+CD25- T cells and HLA-DR4+ BMDCs, we observed a dose-dependent response to citTNC-T4 pulsed BMDCs (fig. 5A), indicating that the citTNC22 peptide is a biologically processed antigen and thus a candidate autoantigen in RA.

**Figure 5.**
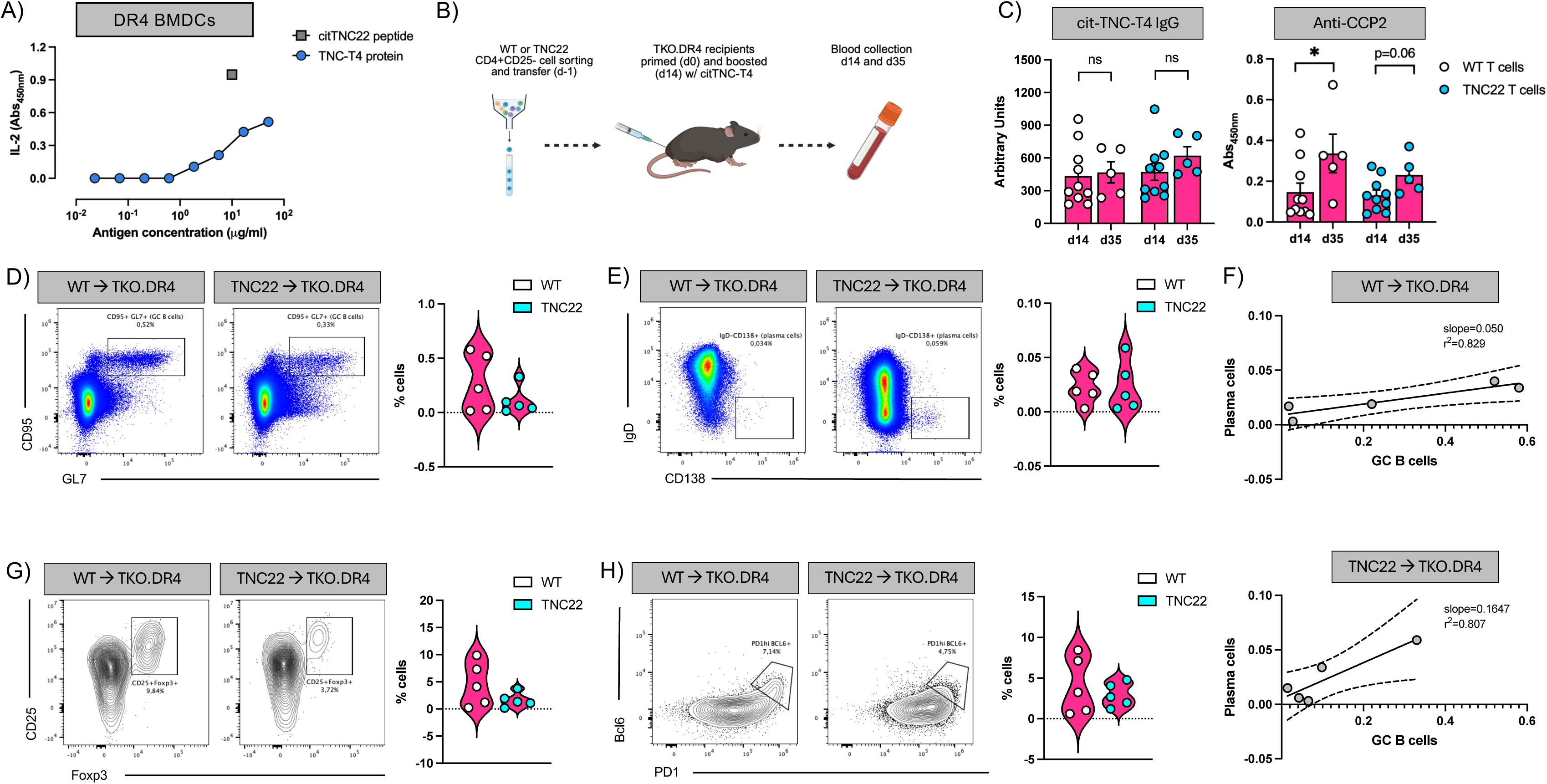
Reactivity of citTNC22 T cells in the context of HLA-DR4 and effect on antibody production. A) Reactivity of TNC22 CD4^+^CD25^-^ T cells in vitro towards PAD-treated TNC-T4 protein fragment and citTNC22 peptide in the presence of DR4^+^ BMDCs. B) Schematic representation of cell transfer, immunization and sample collection for antibody titer analysis. C) citTNC-T4 specific IgG and anti-CCP2 levels at days 14 and 35 post initial immunization (1:25 plasma dilution). D) Germinal center (GC; CD95^+^GL7^+^) B cells in day 14 iLNs of TKO.DR4 recipients from (B). E) Plasma cells (IgD^-^CD138^+^) cells in day 14 iLNs of TKO.DR4 recipients from (B). F) Correlation between GC B cells and plasma cells in day 14 iLNs of TKO.DR4 recipients from (B). G) CD25^+^Foxp3^+^ T cells in day 14 iLNs of TKO.DR4 recipients from (B). H) Tfh (PD1^hi^Bcl6^+^) cells in day 14 iLNs of TKO.DR4 recipients from (B).

We initially showed that some HLA-DR4 transgenic mice harbouring polyclonal T cells can mount a response to citTNC22, and that anti-citrulline antibody titres develop in these mice upon antigen priming and boosting (fig. 1). To circumvent this variable polyclonal T cell reactivity, we sorted CD4+CD25- T cells from hTCR-tg TNC22.TKO mice and their wild-type littermates (WT.H-2^b^) and adoptively transferred them to TKO.DR4 recipient mice. We focused on a cell transfer setting where T cells were educated in a H-2^b^ background, thus ensuring that neither polyclonal (WT.H-2^b^) nor hTCR-tg T cells (TNC22.TKO) had previously been exposed to citTNC22 in the context of HLA-DR4. Mice were then primed and boosted (14 days later) with citTNC-T4 protein and blood was collected at different time points to assess antigen-specific antibody titres (fig. 5B). Both sets of recipients generated an antigen-specific IgG response to the immunized antigen, as well as anti-CCP titres (fig. 5C), demonstrating that hTCR-tg cells can support antigen-specific B cell activation and class-switch. Although recipients of TNC22 T cells showed an apparent reduced frequency of germinal centre (GC) B cells compared to recipients of polyclonal wild-type T cells, the correlation between GC B cells and plasma cells was similar between the two groups (fig. 5D-F), as was the frequency of follicular helper T cells (fig. 5G), in line with the similar antibody levels observed (fig. 5C). Interestingly, hTCR-tg T cells generated less Treg cells than their wild-type polyclonal counterparts, supporting a stronger reactivity of these transgenic T cells upon antigen encounter (fig. 3D-H). Together with the data presented above (fig. 3 and fig. 5A), this observation indicates that hTCR-tg T cells can recognize their cognate citrullinated antigen after biological processing (i.e. citrullinated TNC-T4), mediating effector functions in vivo including supporting of B cell activation and class-switching of B cells recognizing TNC.

### hTCR-tg mice develop inflammatory arthritis upon articular challenge

In the pursuit of a mouse model of RA that better mimics the human disease, we explored the development of arthritis in TNC22 mice. Previous reports show that TNC is expressed in rodent bones upon mechanical loads[34], which we confirmed in wild type mice undergoing inflammatory arthritis (suppl. fig. 7A). The imaging and staining pattern indicate that TNC is expressed within the inflamed synovium, where local stromal cells (i.e. fibroblasts) are likely responsible for such expression[22]. However, systemic immunizations with citTNC22 peptide or with citTNC protein did not give rise to arthritis neither in hTCR-tg mice expressing H-2^b^ nor in double transgenic mice expressing both hTCR and the human HLA-DR4 molecule (TNC22.DR4).

Thus, we investigated whether a local, rather than systemic, antigen challenge would be able to induce inflammatory arthritis. CD4+CD25- hTCR-tg T cells from TNC22.TKO mice were sorted and stimulated with citTNC22 peptide in vitro for 3 days. Activated cells were then transferred into TKO.DR4 recipient mice, followed by systemic immunization with their cognate antigen. Ten days later, a periarticular challenge with antigen was performed in one of the recipient hind ankles (fig. 6A). Arthritis developed in the challenged paw, less than 24h after challenge, covering both the ankle and knuckles of the paw (fig. 6B-C and suppl. fig. 7B). Taken together, the establishment of this novel humanized mouse strain expressing a patient-derived TCR reactive to a PTM autoantigen in the context of a disease-relevant HLA-DR allele will allow further studies of how anti-citrulline T cell reactivity originates and how it is tolerized. This is of importance, since such endeavours are difficult to achieve in existing murine arthritis models where anti-citrulline T cell immunity is not as prominent as it is in the human disease.

**Figure 6.**
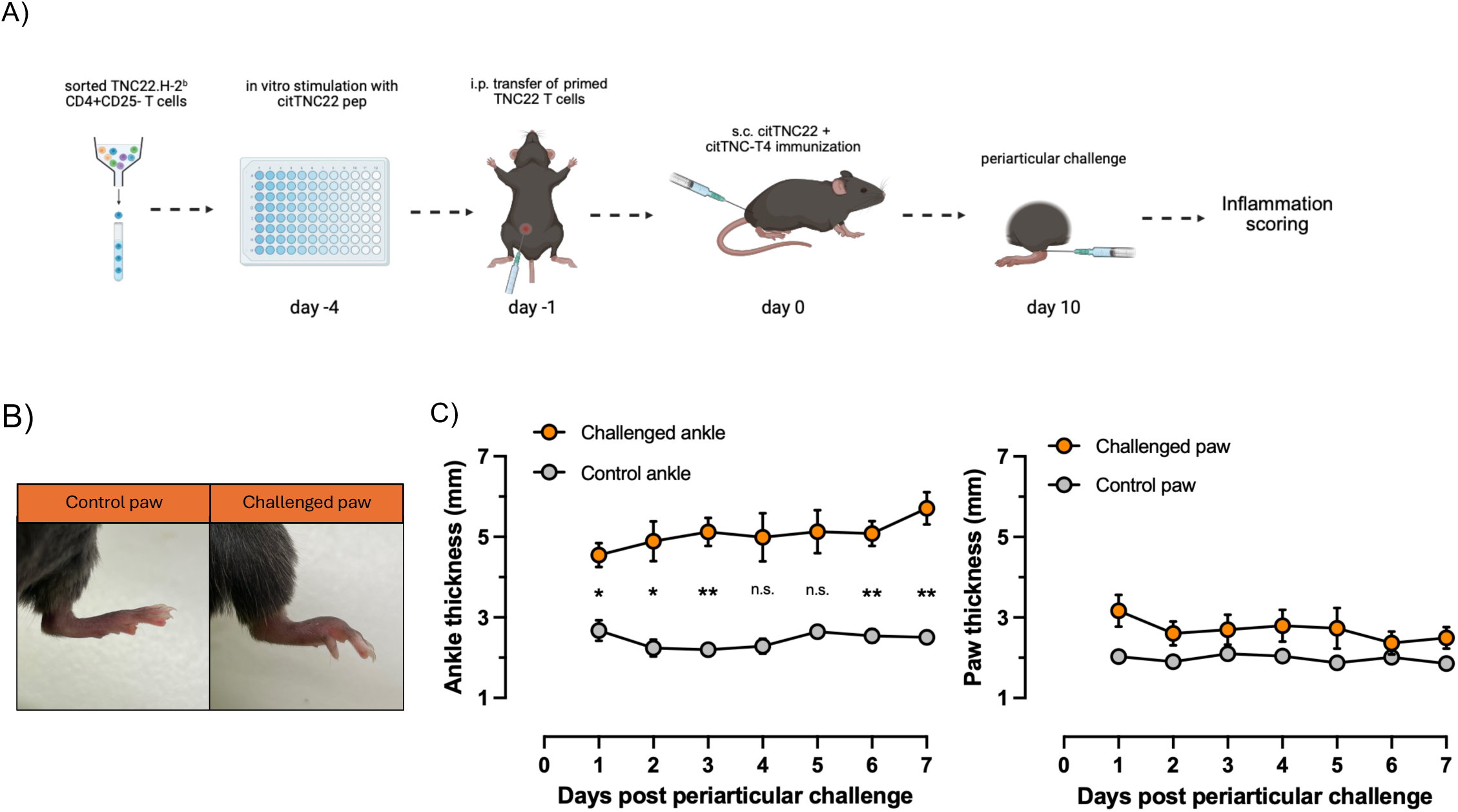
Arthritis development in periarticularly challenged TKO.DR4 mice containing hTCR-tg T cells. A) Schematic representation of arthritis induction protocol in TKO.DR4 mice receiving primed TNC22 CD4^+^CD25^-^ T cells. B) Representation of arthritis phenotype in a control versus challenged paw in the same animal. C) Inflammatory assessment of arthritis measured by thickness of paw and ankle in control and challenged hind limbs. N=4; 2way ANOVA with Tukey’s multiple comparison test.

## Discussion

Human citrulline-specific T cell responses have so far mainly been described towards epitopes identified in proteins previously shown to be prominent targets of RA-specific autoantibodies. Such T cells have been described to be preferentially present in inflamed synovial compartments[11,17], and recently also in the lungs[35] where initiation of RA specific anti-citrulline immunity has been postulated to take place[36]. This is a disconnect to the many animal models of immune-mediated arthritis currently used to study RA pathogenesis and potential therapies[37]. The relative lack of experimental models mimicking the anti-citrulline T- and B-cell reactivity frequently observed in patients is of particular importance when efforts aiming to modulate or eliminate antigen-specific immunity are put forward. Here, we set out to establish a more translational mouse model of RA with a focus on citrulline-specific autoimmunity that is prominent in RA patients by using a patient-derived TCR sequence of known autoreactivity[18,20]. We chose to generate a transgenic mouse line with a chimeric TCR recognizing a citrullinated epitope within tenascin C in the context of HLA-DRB1*04:01. Tenascin C is an autoantigen with potential high relevance to the immune pathology, with T cells recognizing different citrullinated epitopes within this protein being found in both patients with established disease and in individuals at risk of developing RA[17–19].

We demonstrate that wild-type HLA-DR4 mice can mount a polyclonal T cell response to citTNC22 with variable degrees of reactivity, suggesting that tolerance mechanisms towards this antigen may be in place. Our results also support the notion that PTM-specific autoreactivities can be studied in an experimental setting and in an HLA-restricted manner. The development of citTNC22-reactive hTCR-tg mice was intended to investigate the T cell reactivity to this RA-relevant autoantigen in a more controlled setting and provide a tool for future exploration of novel therapies to RA. To our knowledge, this is the first time a humanized mouse strain has been developed to express a patient-derived autoreactive TCR towards a relevant RA antigen and in the context of an HLA-DR allele strongly associated with RA susceptibility. In this initial characterization, we showed that hTCR-tg TNC22 mice develop normally and are fertile. Only in combination with HLA-DR4 (TNC22.DR4) do mature T cells preferentially express the transgenic TCR (suppl. fig. 2G and suppl. fig. 6E), although selective pressures still affect thymic populations and consequently the output to the periphery (fig. 2 and 4). This observation is particularly important in the context of human RA and HLA-DRB1*04:01 individuals, as it suggests that the selection of this autoreactive TCR may result not only from a defect in negative-selection towards the PTM antigen, but also an impaired positive selection of the TCR towards HLA-DR4. Similar observations have been made when re-expressing different hTCRs with the same specificity in the context of type 1 diabetes and HLA-DQ8 expression[38]. Further studies using this hTCR and other RA-associated autoreactive hTCRs will be essential to better answer these questions.

hTCR-tg mice successfully develop CD4+ T cells that show specificity to the citTNC22 peptide without cross-reacting with its native non-modified form. This reactivity is restricted to the presentation of the citrullinated peptide by the human HLA-DR4 molecule, with no reactivity to other disease-relevant citrullinated antigens. Nevertheless, the development of a higher-than-average frequency of Foxp3+ T cells in these mice suggests that this TCR may be promiscuous to other (yet unidentified) antigens of murine origin. Because TNC22-tg CD4^+^ T cells are unable to respond to either the native or citrullinated TNC22 peptides when presented by the murine H-2^b^ allele (fig. 3), we anticipate that the increase of Foxp3+ T cells in these mice is independent of T cell reactivity towards TNC22. This phenotype was partially observed in an HLA-DR4 background, although at a significantly lower level. Hence, in order to mitigate any eventual by-stander regulation by these Foxp3^+^ T cells, we have performed our in vivo experiments using sorted CD4^+^CD25^-^ T cells from TNC22.H-2^b^ mice and transferring them to HLA-DR4+ recipients, where CD25 was used as a proxy to exclude regulatory T cells. Importantly, hTCR-tg T cells demonstrated a capacity to support B cell activation, Ig class-switch and antigen-specific antibody production (fig. 5). This is an essential feature in RA, as CD4+ T cells are important mediators behind the generation and continuous refinement of ACPA in the human disease (discussed in ref: [3]).

In addition to generating a model relevant to study anti-citrulline autoimmunity, we also explored whether humanized HLA-DR4 mice harbouring hTCR-tg TNC22 CD4+ T cells could develop arthritis. Double transgenic TNC22.DR4 mice failed to develop any macroscopic signs of inflammatory arthritis either spontaneously or after systemic immunization. Although demonstrating full capacity to respond to their cognate antigen in vivo, hTCR-tg CD4+ T cells may never see their antigen in naive conditions, since TNC is poorly expressed at steady state in the adult mouse[21]. According to this line of thought, it is often considered that a second event is required to mediate the targeting of the joints by autoreactive B- and T-cells[39]. One of such events could be a direct physical trauma in or around the joints. By mimicking a secondary hit with a periarticular antigen challenge in our hTCR-tg mice, we triggered a fast-forwarded process of joint inflammation. The detailed characterization of this disease model is beyond the scope of this report. Conventionally, methylated-BSA [40] and aggregated chicken ovalbumin [41] are used as cationic antigens capable of being retained in the joint and thus elicit an immune response in pre-exposed animals when intraarticularly challenged with the antigen. Importantly, despite the observed disease course being similar to that described for antigen-induced arthritis in mice, the model here presented sets itself apart from these more traditional approaches [40]. First, the antigen tenascin C is an autologous antigen, expressed in the joint under inflammatory conditions (suppl. fig. 7A), and thus a relevant target for immune-mediated inflammatory arthritis, possibly supporting the continuous activation of local autoreactive T cells. Secondly, citrullinated-tenascin C constitutes a relevant autoantigen in RA, as evidenced by the presence of antigen-specific T cells recognizing distinct TNC epitopes in RA patients as well as individuals at risk of developing RA [17–19]. Finally, the model here presented uses a well-characterized patient-derived autoreactive TCR from a highly disease-associated HLA-DRB1 allele, supporting the relevance of this model in the study of cellular and molecular mechanisms involved in human RA, as well as in the development of antigen-specific therapies targeting citrullinated TNC. Moving forward, we intend to further explore this disease model by including multiple patient hTCRs, a combination of hTCRs and citrulline-reactive autoantibodies (ACPA), engineered B cells expressing patient-derived citrulline-reactive B cell receptors, or a combination of all the above.

The current results demonstrate the potential of these novel hTCR-tg mice as a model to study citrulline-reactive T cells relevant to RA. In future endeavours, we intend using this model to investigate the interplay between autoreactive T and B cells targeting citrullinated antigens, at the same time we explore pre-clinical T cell tolerance approaches with translational capacity.

## Acknowledgements and affiliations

We would like to thank Genadiy Kozhukh for the help in the purification process of the TNC-T4 protein fragment used in parts of this study. We also thank Amin Ravaei for helping us elaborating a reply to a reviewer’s comment. This research project was partly supported by a research grant from *Pfizer Inc.* F.W. has received consulting fees from SmartCella Solutions and Chiesi Pharmaceuticals Inc outside of the scope of the present study.

V.R. received funding from the *Swedish Research Council* (2022-01679, 2022), *Rheumatiker Förbundet* (R-995666, 2023) and *Konung Gustaf V:s 80-årsfond Foundation* (FAI-2022-0901, 2022). B.R. received funding from the *Konung Gustaf V:s 80-årsfond Foundation* (FAI-2022-0870, 2022; FAI-2023-1047, 2023), *Ulla and Gustaf af Uggla Foundation* (FS-2022-0011, 2022) and *Professor Nanna Svartz Foundation* (2022-00459, 2022). M.S. was supported by the Karolinska Institutet KID grant for doctoral studies (2022-00930, 2022).

Part of the data in this manuscript has been orally presented at the 7^th^ European Conference of Immunology (2024) and the published abstract can be found at DOI: 10.1002/eji.202470200.

## Contributions

MSc, MA, MSi, MH, VM and BRa designed experiments, analysed data and wrote the manuscript along with AD, BRe, FW, AW, LK and AL. DVP and AL designed the constructs and helped generating the hTCR-tg mouse strain. MSc and BRa conducted all animal experiments and handling of samples.

## Key Messages

`o` **What is already known on this topic** – Although rare and difficult to study, autoreactive T cells are associated with development of RA and maturation of ACPA. Current animal models fail to mimic these immune responses in humans further limiting studies dissecting citrulline autoimmunity and how to interfere with it. The establishment of a model better resembling the human disease would allow the study of anti-citrulline immune responses characteristic of RA, while simultaneously provide a platform for testing and developing novel antigen-specific therapies.
`o` **What this study adds** – We have established a humanized mouse model whose CD4+ T cells express a patient-derived TCR recognizing an autoantigen in the context of a shared epitope allele (HLA-DRB1*04:01). The generated T cells are functional and support B cell activation and differentiation, and hence recapitulates several features of RA.
`o` **How this study might affect research, practice or policy** – This humanized mouse strain establishes the first animal model with exclusive anti-citrulline T cell immunity, opening a series of future research opportunities, including break-of-tolerance mechanisms. Moreover, this new model will provide a platform to enable the development of therapies aimed at eliminating or re-regulating antigen-specific T cell immunity associated with RA.

