## Supplementary figures for "A new humanized TCR transgenic mouse model to study citrullinated tenascin C reactive T cells relevant to rheumatoid arthritis"

#### Supplementary Figure 1

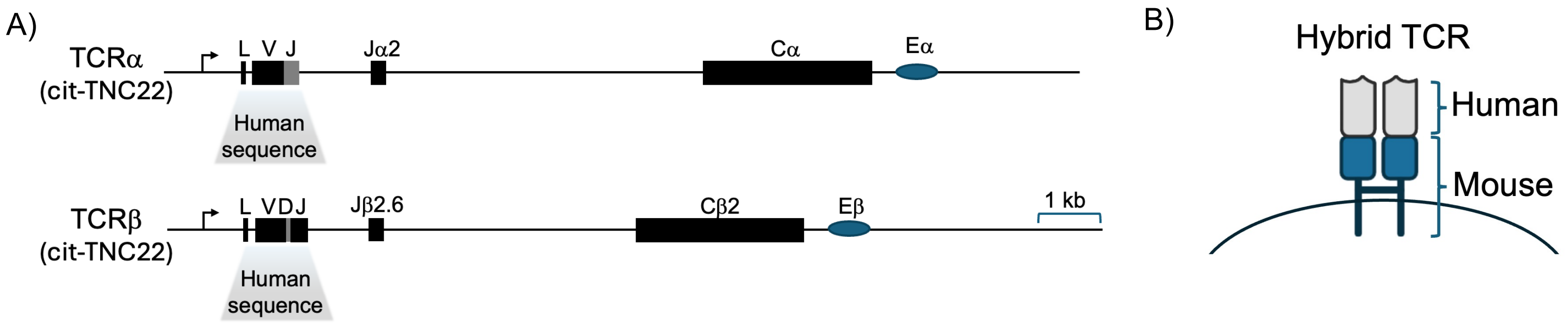

#### Supplementary Figure 2

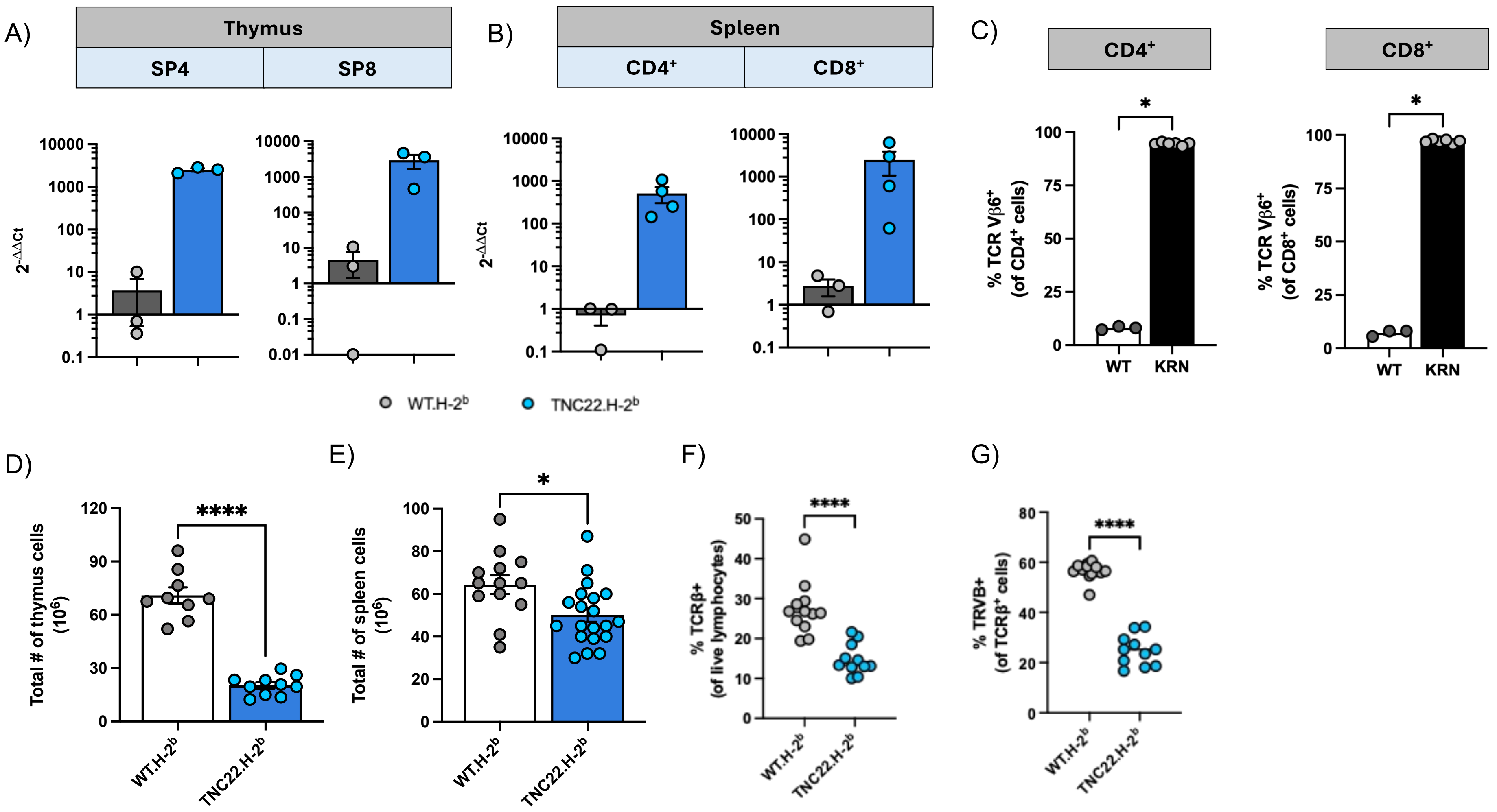

### Supplementary Figure 3

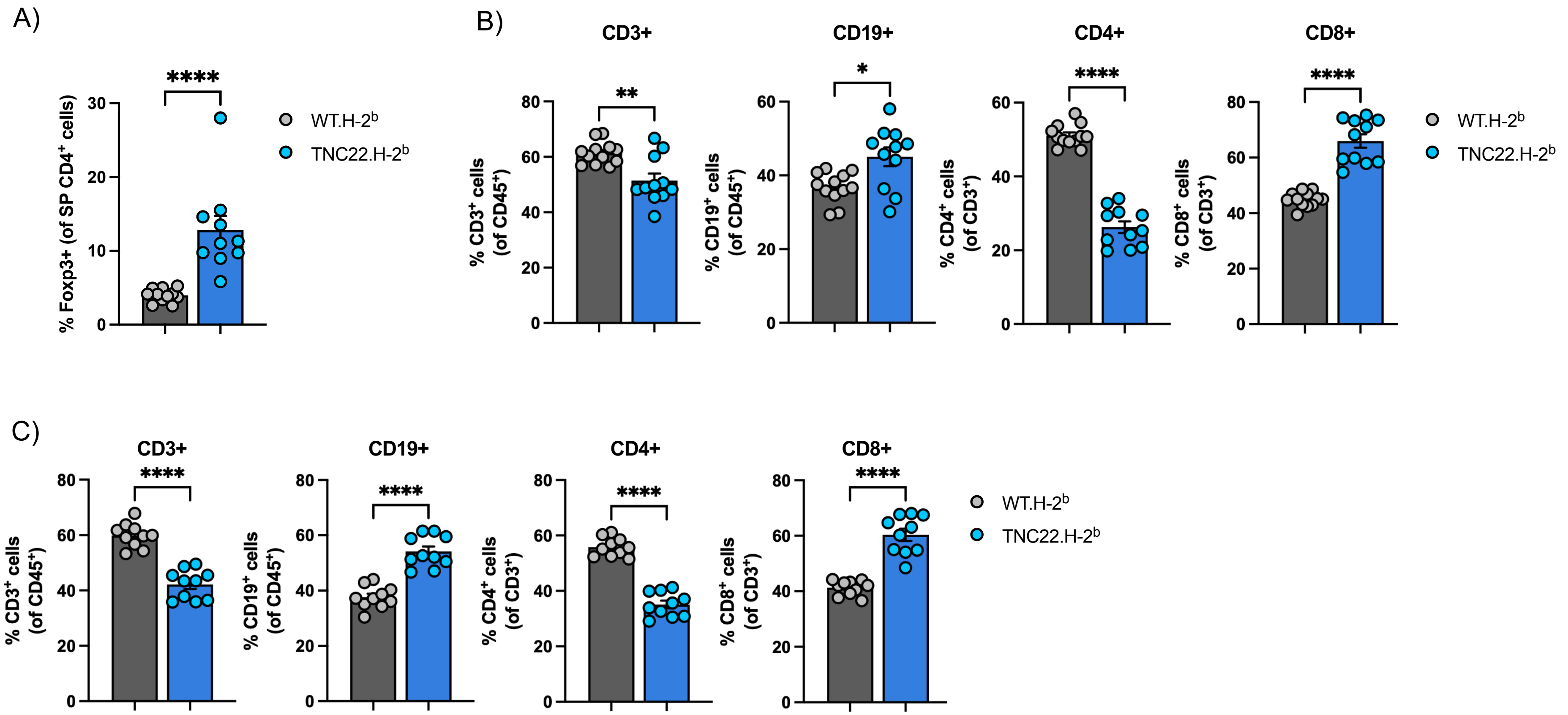

#### Supplementary Figure 4

A)

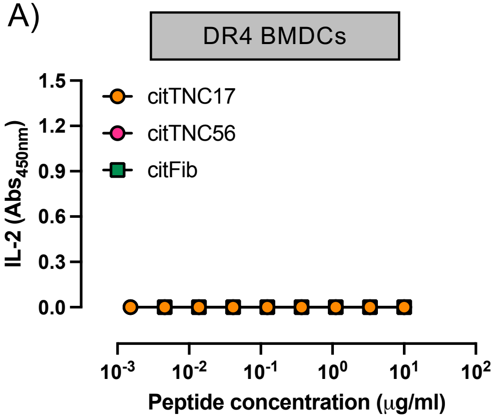


B)

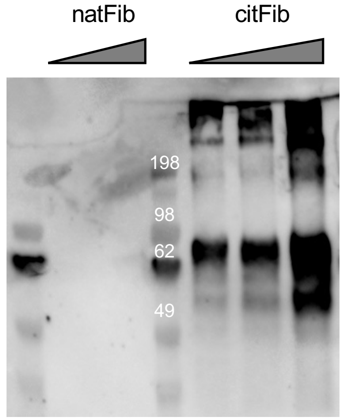

Supplementary Figure 5

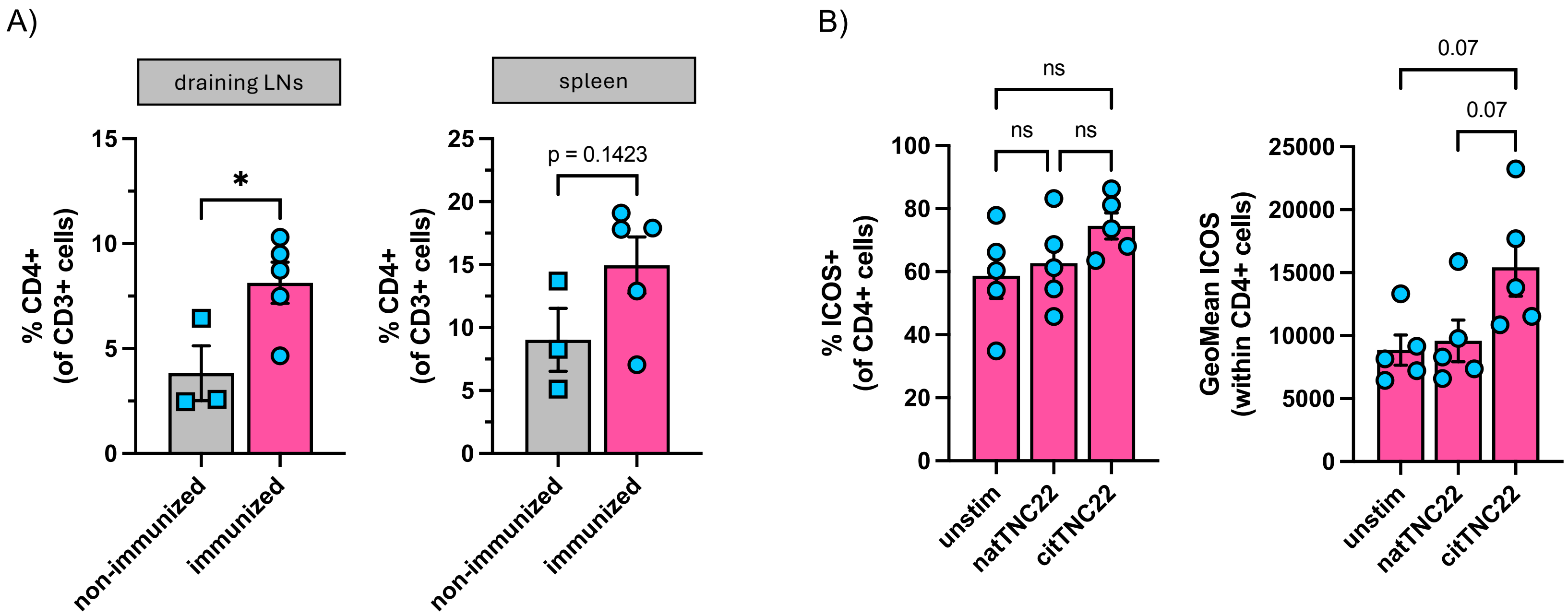

Supplementary Figure 6

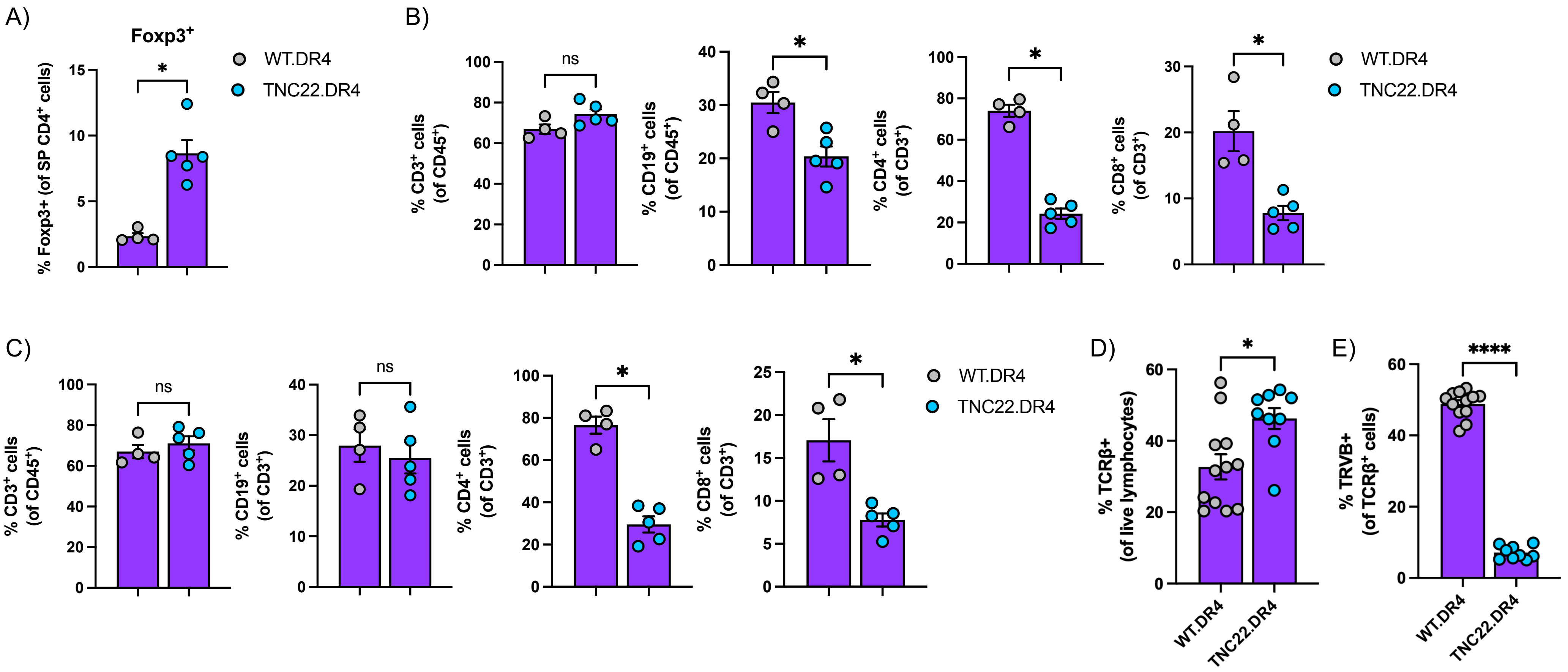

### Supplementary Figure 7

A)

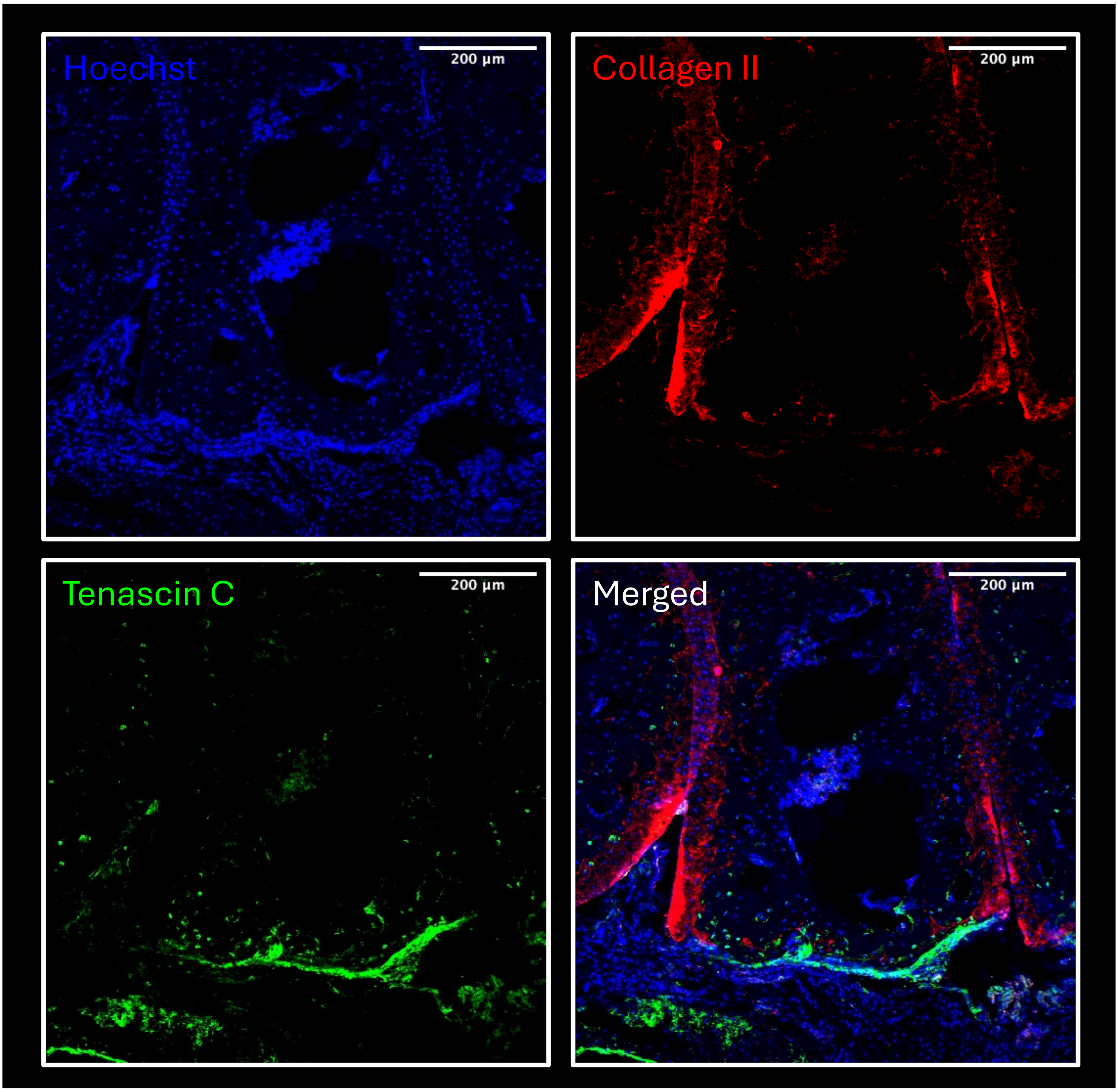


B)

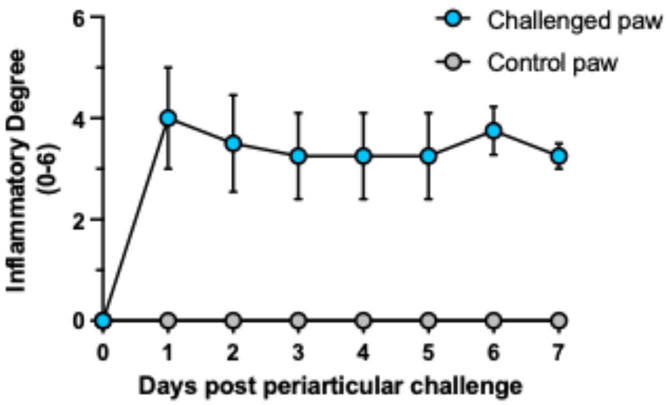
