## Supplementary tables for "A new humanized TCR transgenic mouse model to study citrullinated tenascin C reactive T cells relevant to rheumatoid arthritis"

*Supplementary Table 1: Antibodies anti-mouse used for first surface staining (cocktail #1)*

| **Fluorochrome** | **Marker** | **Clone** | **Host** | **Dilution** | **Supplier** |
|---|---|---|---|---|---|
| BUV563 | HLA-DR | G46-6 | mus IgG2a | 200x | BD Biosciences |
| BV711 | CD8 | 53-6.7 | rat IgG2a | 500x | BioLegend |
| BV750 | CD4 | GK1.5 | rat IgG2b | 500x | Biolegend |
| RB613 | Ly6C | AL-21 | rat IgM | 100x | BD Biosciences |
| RB744 | I-A/I-E | M5/114.15.2 | rat IgG2b | 200x | BD Biosciences |
| PE | TCR Vb 2 | B20.6 | rat IgG2a | 200x | BioLegend |
| PE | TCR Vb 5.1-5.2 | MR9-4 | mus IgG1 | 200x | BioLegend |
| PE | TCR Vb 6 | RR4-7 | rat IgG2b | 200x | BioLegend |
| PE | TCR Vb 7 | TR310 | rat IgG2b | 200x | BioLegend |
| PE | TCR Vb 8.1/8.2 | MR5-2 | mus IgG2a | 200x | BioLegend |
| PE | TCR Vb 8.3 | 1B3.3 | ham IgG | 200x | BioLegend |
| PE | TCR Vb 9 | MR10-2 | mus IgG1 | 200x | BioLegend |
| PE | TCR Vb 11 | RR3-15 | rat IgG2b | 200x | BioLegend |
| PE | TCR Vb 12 | MR11-1 | mus IgG1 | 200x | BioLegend |
| PE | TCR Vb 13 | MR12-4 | mus IgG1 | 200x | BioLegend |
| PE-Cy7 | CD11c | N418 | ham IgG | 200x | BioLegend |
| AF700 | NK1.1 | PK136 | mus IgG2a | 100x | BD Biosciences |

*Supplementary Table 2: Antibodies anti-mouse used for second surface staining (cocktail #2)*

| Fluorochrome | Marker | Clone | Host | Dilution | Supplier |
|---|---|---|---|---|---|
| BUV395 | CD45 | 30-F11 | rat IgG2b | 400x | BD Biosciences |
| BUV737 | CD44 | IM7 | rat IgG2b | 400x | BD Biosciences |
| BV421 | CD25 | PC61 | rat IgG1 | 500x | BioLegend |
| BV480 | CD3e | 145-2C11 | ham IgG1 | 500x | BD Biosciences |
| BV510 | CD11b | M1/70 | rat IgG2b | 500x | BioLegend |
| BV605 | CD62L | MEL-14 | rat IgG2a | 500x | BioLegend |
| BV650 | Ly6G | 1A8 | rat IgG2a | 500x | BioLegend |
| FITC | CD80 | 16-10A1 | ham IgG | 500x | BioLegend |
| FITC | CD86 | GL-1 | rat IgG2a | 500x | BioLegend |
| BB700 | PD1 | 29F.1A12 | rat IgG2a | 100x | BioLegend |
| PE-CF594 | B220 | RA3-6B2 | rat IgG2a | 2000x | BD Biosciences |
| PE-CF594 | CD19 | 1D3 | rat IgG2a | 2000x | BD Biosciences |
| PE-Cy5 | CD69 | H1.2F3 | ham IgG1 | 500x | BioLegend |
| APC | IgD | 11-26c.2a | rat IgG2a | 2000x | BD Biosciences |
