## Supplementary figure legends for "A new humanized TCR transgenic mouse model to study citrullinated tenascin C reactive T cells relevant to rheumatoid arthritis"

**Supplementary Figure 1 – hTCR transgenic construct.**

A-B) Schematic of chimeric mouse/human TCR construct used for the generation of TNC22 mice.

**Supplementary Figure 2 – expression of the transgenic hTCR in TNC22 mice**

A) qPCR from SP4 and SP8 cells of TNC22.H-$2^b$ mice.

B) qPCR from CD4+ and CD8+ splenic cells from TNC22.H-$2^b$ mice.

C) Expression of transgenic Vβ TCR in splenic T cells from KRN transgenic mice, developed using the same backbone plasmid of TNC22 mice. Mann-Whitney test. * - $p<0.05$

D) Total number of thymocytes in TNC22.H-$2^b$ and WT.H-$2^b$ littermate mice. Unpaired t-test. **** - $p<0.0001$

E) Total number of splenocytes in TNC22.H-2b and WT.H-2b littermate mice. Unpaired t-test. * - $p<0.05$

F) Frequency of TCRb+ cells in TNC22.H-$2^b$ and WT.H-$2^b$ littermate mice. Unpaired t-test. **** - $p<0.0001$

G) Frequency of murine TRBV+ cells in TNC22.H-$2^b$ and WT.H-$2^b$ littermate mice. Unpaired t-test. **** - $p<0.0001$

**Supplementary Figure 3 – hTCR-tg T cell phenotypes in skin and mucosa draining lymph nodes in a H-2$^b$ background.**

A) Frequency of Foxp3+ CD4+ T cells in thymi of TNC22.H-2$^b$ and WT.H-2$^b$ littermates. Unpaired t-test. **** - p<0.0001

B-C) Characteristics of T cells originating from iLN (B) and mLN (C) of TNC22.H-2$^b$ and WT.H-2$^b$ littermates. Unpaired t-test. * - p<0.05; ** - p<0.01; **** - p<0.0001

**Supplementary Figure 4 – Reactivity of TNC22 T cells to other citrullinated peptides than TNC22.**

A) IL-2 production by TNC22 CD4+CD25- T cells to citrullinated peptides within TNC and citrullinated fibrinogen, in the presence of DR4+ BMDCs.

B) Confirmation of in vitro citrullination of PAD-treated fibrinogen, by western blot.

**Supplementary Figure 5 – TNC22 T cells expansion upon antigen encounter in vivo.**

A) Donor TNC22 CD4+ T cells expand in citTNC22-immunized recipient DR4+ mice.

B) Upregulation of ICOS on transferred TNC22 CD4+ T cells after antigen recall of splenocytes isolated from citTNC22-immunized DR4+ recipient mice.

**Supplementary Figure 6 – hTCR-tg T cell phenotypes in skin and mucosa draining lymph nodes in an HLA-DR4 background.**

A) Thymus frequency of Foxp3+ CD4+ T cells in TNC22.DR4 mice, compared to WT.DR4 littermates. *Mann-Whitney unpaired test*. * – $p<0.05$

B-C) iLN (B) and mLN (C) characteristics of T cells in TNC22.DR4 mice and WT.DR4 littermates. *Mann-Whitney unpaired test.* * – $p<0.05$; n.s. – not significant

D) Frequency of TCRb+ cells in TNC22.DR4 and WT.DR4 littermate mice. Unpaired t-test. * – $p<0.05$

E) Frequency of murine TRBV+ cells in TNC22.DR4 and WT.DR4 littermate mice. Unpaired t-test. **** – $p<0.0001$

**Supplementary Figure 7 – TNC is an antigen present within the inflamed synovium.**

A) Identification of tenascin C (green), type II collagen (red) and cell nucleus (blue) in arthritic paw of a CAIA Balb/c mouse, 14 days post disease induction (10x objective).

B) Qualitative scoring of arthritis in HLA-DR4 recipients of hTCR-tg TNC22 T cells after periarticular challenge.
